## Appendix 1 for "Novel methods for epistasis detection in genome-wide association studies"

### S2 Appendix. Simulation results

### 1 First scenario: synergistic only effects

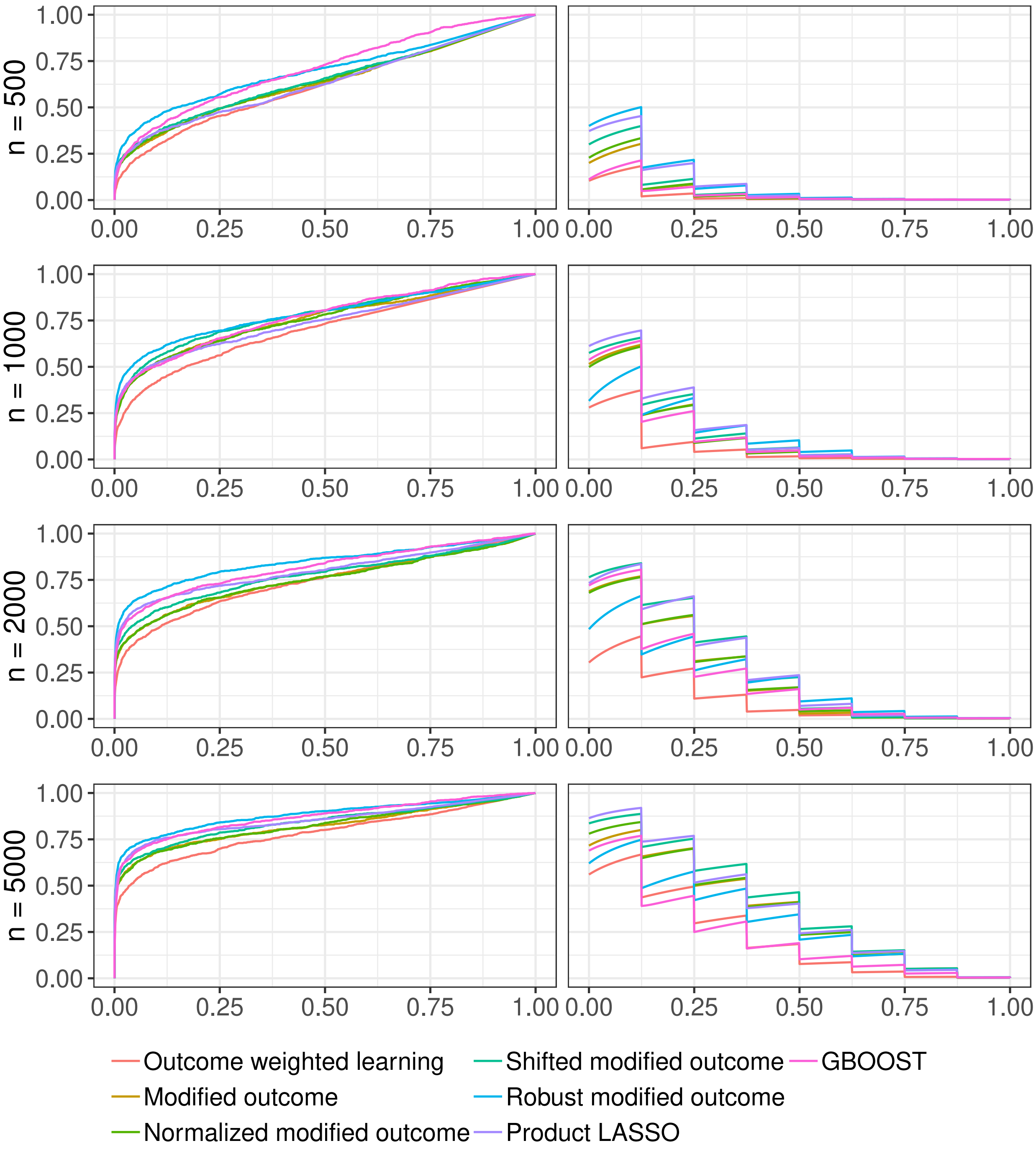


**Fig A.** Average ROC (left column) and PR (right column) curves for the first scenario

**Table A.** Average ROC and PR AUCs for the first scenario

| Method | PR | ROC |
|---|---|---|
| n =500 | | |
| GBOOST | 0.0362 | 0.7075 |
| Modified outcome | 0.0468 | 0.6747 |
| Robust modified outcome | 0.0973 | 0.7414 |
| Normalized modified outcome | 0.0512 | 0.6754 |
| Shifted modified outcome | 0.0644 | 0.6794 |
| Outcome weighted learning | 0.0254 | 0.6282 |
| Product LASSO | 0.0895 | 0.6514 |
| n =1000 | | |
| GBOOST | 0.1270 | 0.7688 |
| Modified outcome | 0.1284 | 0.7131 |
| Robust modified outcome | 0.1302 | 0.7434 |
| Normalized modified outcome | 0.1255 | 0.7120 |
| Shifted modified outcome | 0.1470 | 0.7224 |
| Outcome weighted learning | 0.0613 | 0.6764 |
| Product LASSO | 0.1619 | 0.7032 |
| n =2000 | | |
| GBOOST | 0.2103 | 0.8169 |
| Modified outcome | 0.2252 | 0.7512 |
| Robust modified outcome | 0.2070 | 0.8449 |
| Normalized modified outcome | 0.2266 | 0.7501 |
| Shifted modified outcome | 0.2704 | 0.7753 |
| Outcome weighted learning | 0.1045 | 0.7394 |
| Product LASSO | 0.2711 | 0.7989 |
| n =5000 | | |
| GBOOST | 0.2276 | 0.8697 |
| Modified outcome | 0.3512 | 0.8218 |
| Robust modified outcome | 0.3011 | 0.8818 |
| Normalized modified outcome | 0.3548 | 0.8248 |
| Shifted modified outcome | 0.3907 | 0.8423 |
| Outcome weighted learning | 0.2139 | 0.7847 |
| Product LASSO | 0.3779 | 0.8546 |

### 2 Second scenario: partial overlap between synergistic and marginal effects

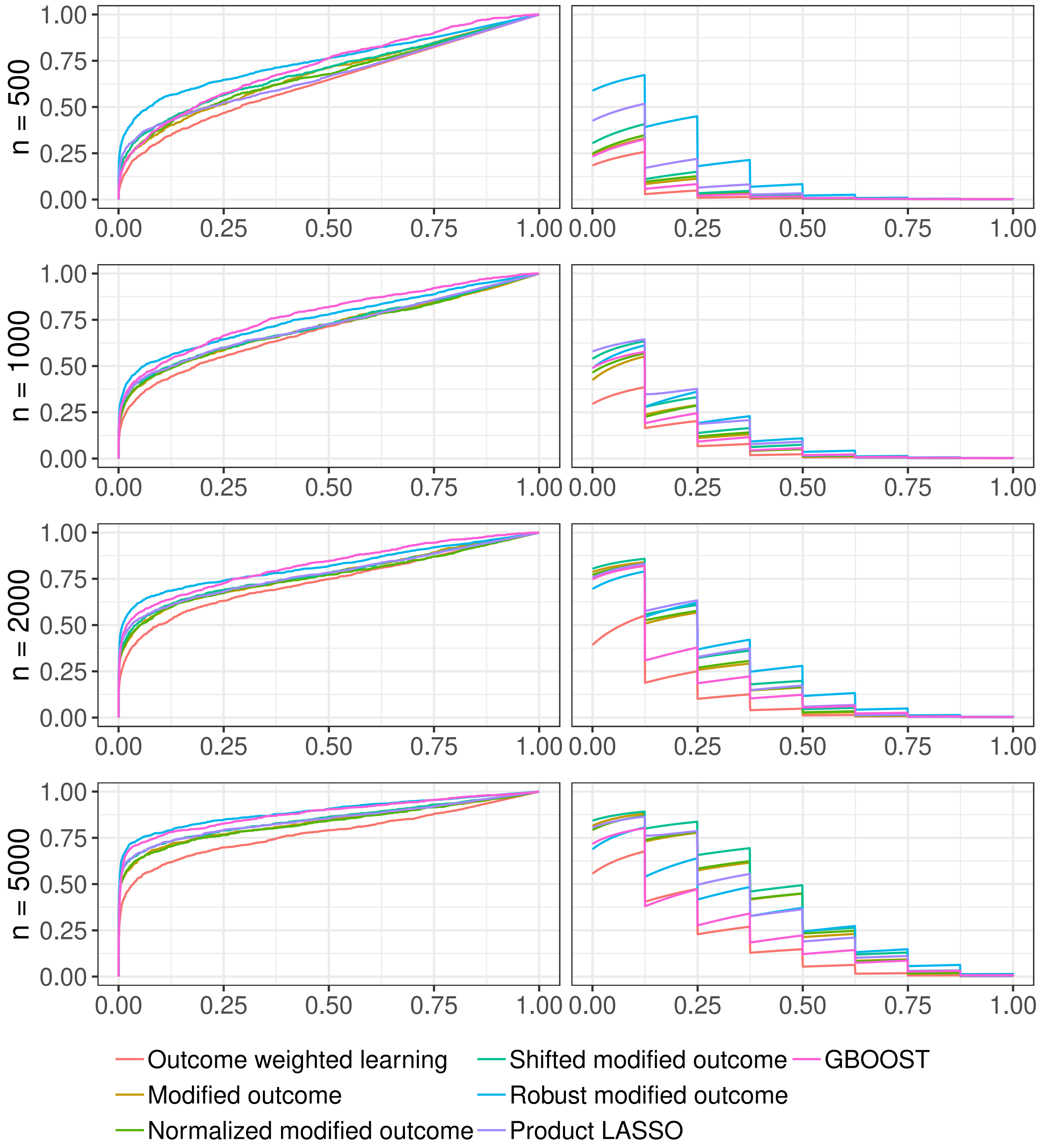


**Fig B.** Average ROC (left column) and PR (right column) curves for the second scenario

**Table B.** Average ROC and PR AUCs for the second scenario

| Method | PR | ROC |
|---|---|---|
| n =500 | | |
| GBOOST | 0.0516 | 0.7186 |
| Modified outcome | 0.0563 | 0.6750 |
| Robust modified outcome | 0.1716 | 0.7502 |
| Normalized modified outcome | 0.0590 | 0.6713 |
| Shifted modified outcome | 0.0712 | 0.6918 |
| Outcome weighted learning | 0.0367 | 0.6345 |
| Product LASSO | 0.0994 | 0.6659 |
| n =1000 | | |
| GBOOST | 0.1190 | 0.7773 |
| Modified outcome | 0.1195 | 0.7092 |
| Robust modified outcome | 0.1574 | 0.7601 |
| Normalized modified outcome | 0.1233 | 0.7080 |
| Shifted modified outcome | 0.1443 | 0.7160 |
| Outcome weighted learning | 0.0805 | 0.6923 |
| Product LASSO | 0.1609 | 0.7170 |
| n =2000 | | |
| GBOOST | 0.1933 | 0.8226 |
| Modified outcome | 0.2294 | 0.7708 |
| Robust modified outcome | 0.2732 | 0.8183 |
| Normalized modified outcome | 0.2321 | 0.7623 |
| Shifted modified outcome | 0.2532 | 0.7753 |
| Outcome weighted learning | 0.1114 | 0.7360 |
| Product LASSO | 0.2507 | 0.7762 |
| n =5000 | | |
| GBOOST | 02454 | 0.8821 |
| Modified outcome | 0.3718 | 0.8344 |
| Robust modified outcome | 0.3286 | 0.8916 |
| Normalized modified outcome | 0.3739 | 0.8309 |
| Shifted modified outcome | 0.4079 | 0.8487 |
| Outcome weighted learning | 0.1930 | 0.7769 |
| Product LASSO | 0.3537 | 0.8467 |

### 3 Third scenario: partial overlap between synergistic and quadratic effects

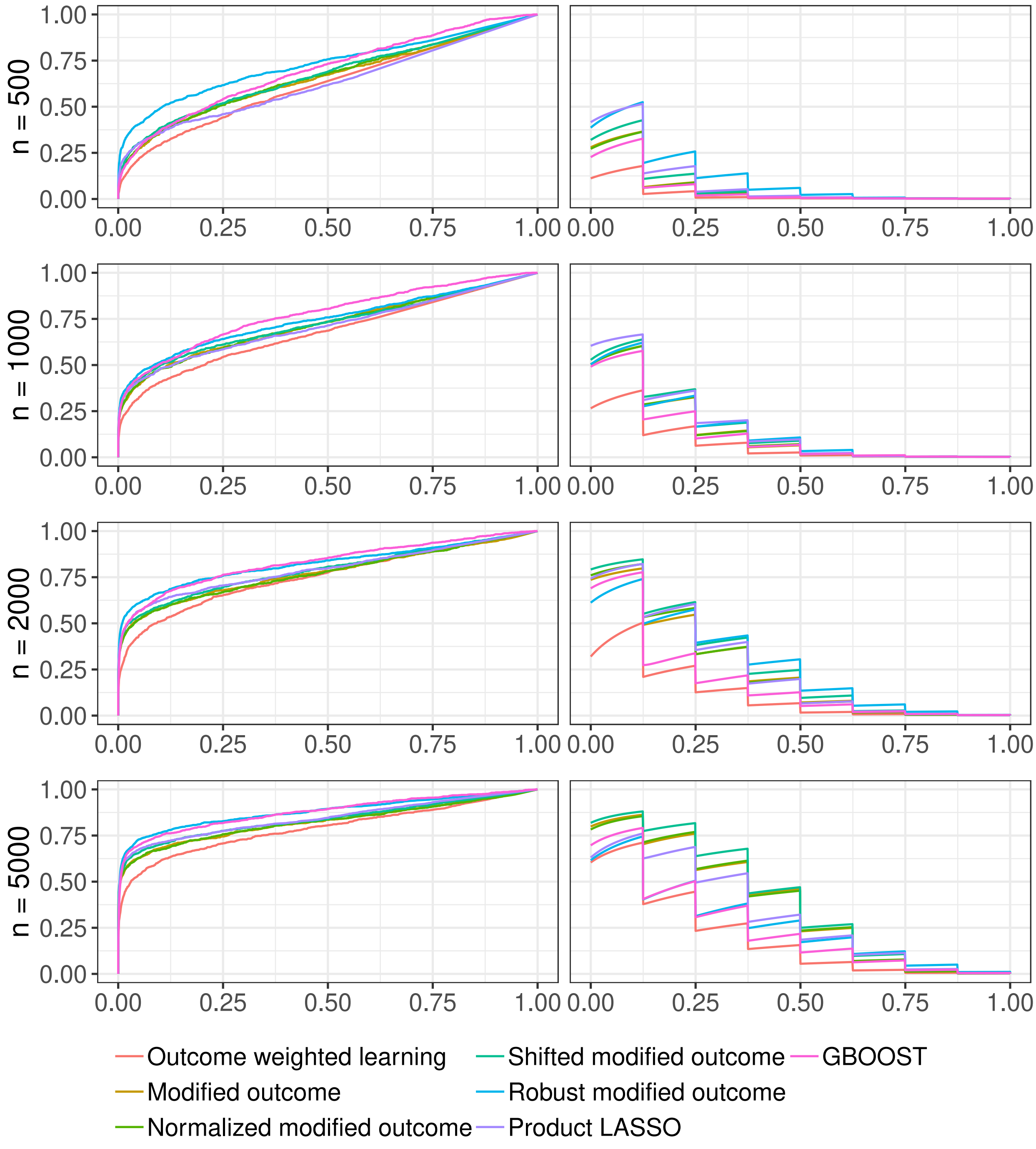


**Fig C.** Average ROC (left column) and PR (right column) curves for the third scenario

**Table C.** Average ROC and PR AUCs for the third scenario

| Method | PR | ROC |
|---|---|---|
| n =500 | | |
| GBOOST | 0.050 | 0.6970 |
| Modified outcome | 0.0570 | 0.6559 |
| Robust modified outcome | 0.1148 | 0.7296 |
| Normalized modified outcome | 0.0569 | 0.6627 |
| Shifted modified outcome | 0.0714 | 0.6703 |
| Outcome weighted learning | 0.0260 | 0.6233 |
| Product LASSO | 0.0889 | 0.6282 |
| n =1000 | | |
| GBOOST | 0.1228 | 0.7746 |
| Modified outcome | 0.1362 | 0.7181 |
| Robust modified outcome | 0.1513 | 0.7444 |
| Normalized modified outcome | 0.1373 | 0.7175 |
| Shifted modified outcome | 0.1546 | 0.7226 |
| Outcome weighted learning | 0.0728 | 0.6778 |
| Product LASSO | 0.1620 | 0.7100 |
| n =2000 | | |
| GBOOST | 0.1814 | 0.8307 |
| Modified outcome | 0.2430 | 0.7733 |
| Robust modified outcome | 0.2697 | 0.8235 |
| Normalized modified outcome | 0.2496 | 0.7724 |
| Shifted modified outcome | 0.2737 | 0.7886 |
| Outcome weighted learning | 0.1129 | 0.7535 |
| Product LASSO | 0.2543 | 0.7921 |
| n =5000 | | |
| GBOOST | 0.2467 | 0.8767 |
| Modified outcome | 0.3663 | 0.8241 |
| Robust modified outcome | 0.2660 | 0.8790 |
| Normalized modified outcome | 0.3669 | 0.8236 |
| Shifted modified outcome | 0.3944 | 0.8376 |
| Outcome weighted learning | 0.1965 | 0.7893 |
| Product LASSO | 0.3158 | 0.8439 |

### 4 Fourth scenario: partial overlap between synergistic and quadratic/marginal effects

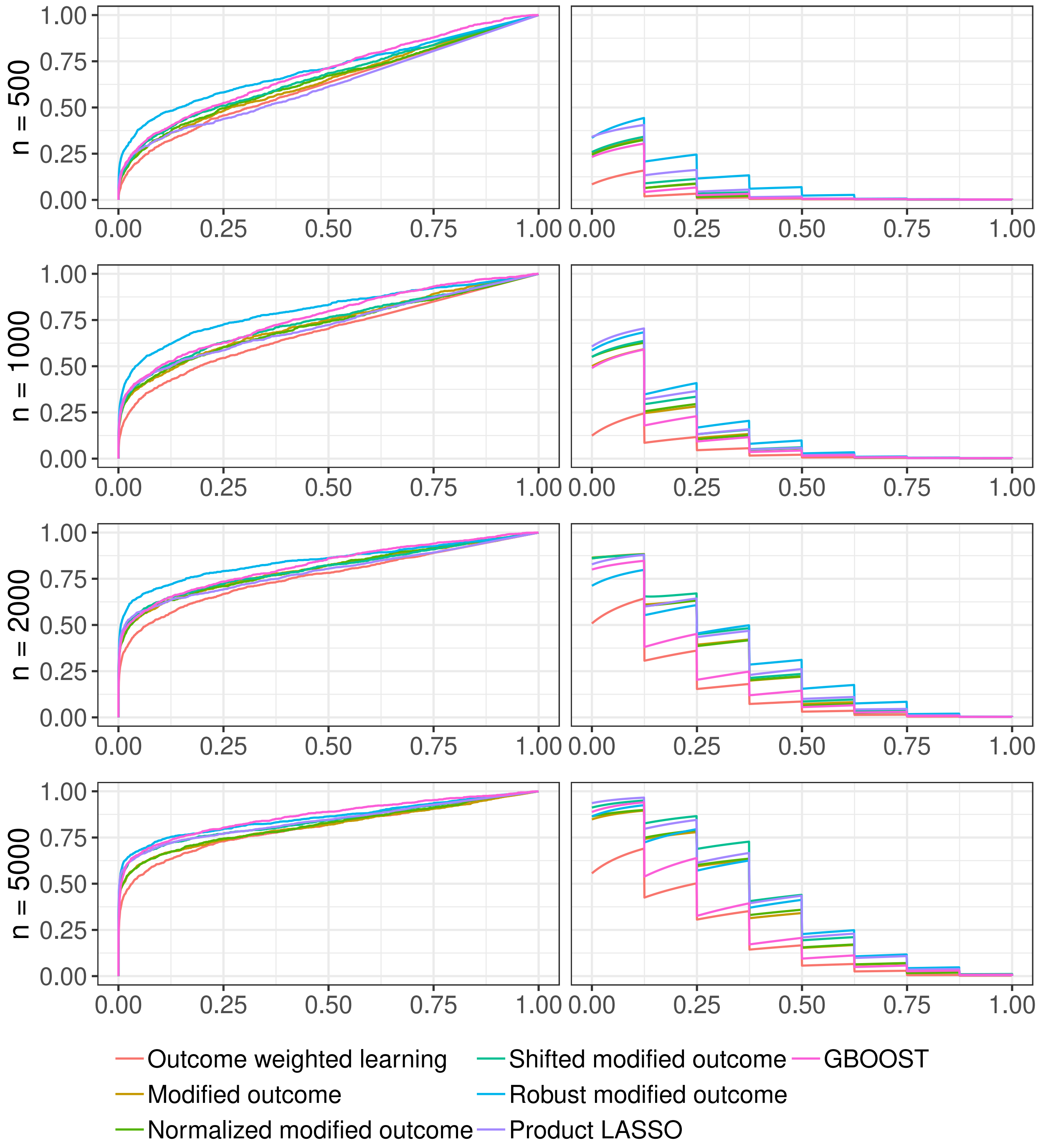


**Fig D.** Average ROC (left column) and PR (right column) curves for the fourth scenario

**Table D.** Average ROC and PR AUCs for the fourth scenario

| Method | PR | ROC |
|---|---|---|
| n =500 | | |
| GBOOST | 0.0479 | 0.6900 |
| Modified outcome | 0.0521 | 0.6427 |
| Robust modified outcome | 0.1066 | 0.7065 |
| Normalized modified outcome | 0.0513 | 0.6460 |
| Shifted modified outcome | 0.0591 | 0.6623 |
| Outcome weighted learning | 0.0227 | 0.6218 |
| Product LASSO | 0.0762 | 0.6174 |
| n =1000 | | |
| GBOOST | 0.1163 | 0.7647 |
| Modified outcome | 0.1283 | 0.7288 |
| Robust modified outcome | 0.1687 | 0.8049 |
| Normalized modified outcome | 0.1338 | 0.7200 |
| Shifted modified outcome | 0.1438 | 0.7388 |
| Outcome weighted learning | 0.0479 | 0.6838 |
| Product LASSO | 0.1554 | 0.7206 |
| n =2000 | | |
| GBOOST | 0.2129 | 0.8237 |
| Modified outcome | 0.2794 | 0.8007 |
| Robust modified outcome | 0.2986 | 0.8478 |
| Normalized modified outcome | 0.2763 | 0.8032 |
| Shifted modified outcome | 0.2960 | 0.8050 |
| Outcome weighted learning | 0.1530 | 0.7641 |
| Product LASSO | 0.2927 | 0.7899 |
| n =5000 | | |
| GBOOST | 0.2823 | 0.8656 |
| Modified outcome | 0.3541 | 0.8127 |
| Robust modified outcome | 0.3823 | 0.8568 |
| Normalized modified outcome | 0.3597 | 0.8175 |
| Shifted modified outcome | 0.4091 | 0.8388 |
| Outcome weighted learning | 0.2106 | 0.8031 |
| Product LASSO | 0.4000 | 0.8399 |
