## Appendix 2 for "Novel methods for epistasis detection in genome-wide association studies"

### S1 Appendix. Genotypic hidden Markov model

In this Appendix, we explicit the transition and emission probabilities for the genotypic hidden Markov model. For that purpose, we start by considering a pair of ordered haplotypes $H^a = \left(H_1^a, \cdots, H_p^a\right) \in \{0,1\}^p$ and $H^b = \left(H_1^b, \cdots, H_p^b\right) \in \{0,1\}^p$. We recall that the two haplotypes correspond to the same positions. The hidden variables $Z^a = \left(Z_1^a, \cdots, Z_p^a,\right)$ and $Z^b = \left(Z_1^b, \cdots, Z_p^b,\right)$ represent cluster memberships. They take discrete values in $\{1, \cdots, K\}^p$. Scheet and Stephens [1] define the clusters as a "(common) combination of alleles at tightly linked SNPs". The underlying hidden Markov models for the two alleles have identical forms. We then focus on the first allele $a$. We follow the notations of [2].

The marginal distribution of the first hidden state can be written as:

$$q_1^{hap}(k) = \alpha_{1,k}, \quad k \in \{1, \cdots, K\}.$$

For $j \in \{2, \cdots, p\}$, the transition matrix $Q_j^{hap}$ is given by:

$$Q_j^{hap}(k'|k) = P(H_j = k'|H_{j-1} = k') = \begin{cases} e^{-r_j} + \left(1 - e^{-r_j}\right) \alpha_{j,k'}, & k' = k \\ \left(1 - e^{-r_j}\right) \alpha_{j,k'}, & k' \neq k \end{cases}.$$

The parameter $r = (r_2, \cdots, r_p)$ can be assimilated to the recombination rate between loci $j-1$ and $j$, although Scheet and Stephens [1] point out the general mismatch between the observed recombination rates and the estimate of $r$. The parameter $\alpha = (\alpha_{j,k})_{(j,k)\in\{1,\cdots p\}\times\{1,\cdots,K\}}$ is the relative frequency of the cluster $k$ in locus $j$.

Conditionally on the latent state $Z_j^{hap} = z_j$, the allele $H_j$ is a Bernoulli random variable, $H_j|Z_j \sim \mathcal{B}(\theta_{j,z_j})$. $\theta_{j,z_j}$ is the frequency of allele 1 in cluster $z_j$ at the position $j$:

$$f_j^{hap} = (h_j; z_j, \theta) = \begin{cases} 1 - \theta_{j,z_j}, & h_j = 0 \\ \theta_{j,z_j}, & h_j = 1 \end{cases}.$$

Under the Hardy-Weinberg equilibrium (HWE), a third hidden Markov model for the unphased genotype can be derived by combining the HMMs of the two alleles $a$ and $b$. The emission states $X = (X_1, \cdots, X_p) \in \{0,1,2\}^p$ are given by the sum of the emission states, $H^a + H^b = \left(H_1^a + H_1^b, \cdots, H_p^a + H_p^b\right)$. Because of the phase indetermination, the latent states are unordered pairs of haplotype latent states, $Z = (\{Z_1^a, Z_1^b\}, \cdots, \{Z_p^a, Z_p^b\})$. Thus, the dimensionality of the latent variable space is $K(K+1)/2$. The different probabilities of the genotype model are computed by considering the two cases: $Z_j^a = Z_j^b$ and $Z_j^a \neq Z_j^b$.

The initial latent state distribution is given by:

$$q_1^{gen}(\{k^a, k^b\}) = \begin{cases} (\alpha_{1,k^a})^2, & k^a = k^b \\ 2\alpha_{1,k^a}\alpha_{1,k^b} & k^a \neq k^b \end{cases}.$$

In a similar fashion, the transition probabilities:

$$Q_j^{gen}(\{\underline{k}^a, \underline{k}^b\}|\{k^a, k^b\}) = \begin{cases} Q_j^{hap}(\underline{k}^a|k^a)Q_j^{hap}(\underline{k}^b|k^b) + Q_j^{hap}(\underline{k}^b|k^a)Q_j^{hap}(\underline{k}^a|k^b), & \underline{k}^a \neq \underline{k}^b \\ Q_j^{hap}(\underline{k}^a|k^a)Q_j^{hap}(\underline{k}^b|k^b), & \text{otherwise} \end{cases}.$$

and, the emission probabilities are

$$f_j(x_j; \{k^a, k^b\}, \theta) = \begin{cases} (1-\theta_{j,k^a})(1-\theta_{j,k^b}), & x_j = 0 \\ \theta_{j,k^a}(1-\theta_{j,k^b}) + (1-\theta_{j,k^a})\theta_{j,k^b}, & x_j = 1 \\ \theta_{j,k^a}\theta_{j,k^b}, & x_j = 2 \end{cases}.$$

For the estimate of the parameters $\nu = (\alpha, r, \theta)$, we use the imputation software fastPHASE [1] which fits the hidden Markov model using an expectation-maximization (EM) algorithm [3]. Its computational complexity is $\mathcal{O}(npK^2)$. The complexity scales linearly for both $p$ and $n$, rendering fastPHASE well-suited for real case-control datasets where the number of SNPs is typically in the hundreds of thousands and the number of samples in the thousands. In practice, as a trade-off between a rich representation of the clusters and the ensuing quadratic complexity, we chose $K = 12$.
